# Translational fusion of terpene synthases enhances metabolic flux by increasing protein stability

**DOI:** 10.1101/2022.11.08.515726

**Authors:** Li Chen Cheah, Lian Liu, Terra Stark, Manuel R. Plan, Bingyin Peng, Zeyu Lu, Gerhard Schenk, Frank Sainsbury, Claudia E. Vickers

## Abstract

The end-to-end fusion of enzymes that catalyse successive steps in a reaction pathway is a metabolic engineering strategy that has been successfully applied in a variety of pathways and is particularly common in terpene bioproduction. Despite its popularity, limited work has been done to interrogate the mechanism of metabolic enhancement from enzyme fusion. We observed a remarkable >110-fold improvement in nerolidol production upon translational fusion of nerolidol synthase (a sesquiterpene synthase) to farnesyl diphosphate synthase. This delivered a titre increase from 29.6 mg/L up to 4.2 g/L nerolidol in a single engineering step. Whole-cell proteomic analysis revealed that nerolidol synthase levels in the fusion strains were greatly elevated compared to the non-fusion control. Similarly, the fusion of nerolidol synthase to non-catalytic domains also produced comparable increases in titre, which coincided with improved enzyme expression. When farnesyl diphosphate synthase was fused to other terpene synthases, we observed more modest improvements in terpene titre (1.9- and 3.8-fold), which corresponds to increases of a similar magnitude in terpene synthase expression. Therefore, increased *in vivo* enzyme levels – resulting from improved expression and/or stability – is likely to be a major driver of catalytic enhancement from enzyme fusion.

## INTRODUCTION

Successive reaction steps in a metabolic pathway are often performed either by multi-domain enzymes or enzymes co-localised in clusters, complexes, or organelles^1–3^. Taking inspiration from nature, numerous synthetic biology tools have been developed to spatially organise and co-localise enzymes in engineered bioproduction pathways. The most straightforward way to artificially co-localise two enzymes is to fuse them end-to-end, usually with a short linker peptide in between. The translational fusion of enzymes that catalyse successive steps in a reaction pathway has been widely adopted as a straightforward, orthogonal, and host-agnostic metabolic engineering strategy. This strategy has proven particularly effective in the case of isoprenoid (terpenoid) production, where the fusion of a prenyl diphosphate synthase (such as farnesyl diphosphate synthase, FPPS) with the corresponding terpene synthase improved the production of diverse products such as patchoulol^4,5^, linalool^6^, sabinene^7^, and sclareol^8^.

The observed enhancements from spatially organised enzymes have been hypothesised to originate from either (i) the probabilistic diffusion of the intermediate metabolite towards the second enzyme (rather than into competing pathways), or (ii) the direct transfer of the intermediate metabolite between enzymes without diffusion into the bulk solution^1,9–13^. Specifically for translationally fused enzymes, the former explanation is favoured in the literature^14,15^; this is supported by a study on an FPPS and patchoulol synthase fusion that found constructs designed with short linker peptides slightly outperforming constructs with long linker peptides^4^. Another study investigating the fusion of geranyl diphosphate synthase (GPPS) with pinene synthase also found a small decrease in pinene titre as the linker length increases^16^. Furthermore, an *in vitro* study with purified enzymes found that fusion constructs of FPPS and epi-aristolochene synthase outperformed a mixture of the two individual enzymes in the coupled reaction^17^. However, this does not explain why the opposite was observed in other terpenoid bioproduction studies^5,6^, and the fact that the performance of enzyme fusions can vary when the order of the two fusion partners is swapped^6,16,18–21^. Despite the popularity of the enzyme fusion approach, limited work has been done to interrogate the mechanism(s) of titre enhancement from enzyme fusion in detail.

In this work, we aimed to investigate the mechanistic basis of titre improvements from isoprenoid pathway enzyme fusion using budding yeast *(Saccharomyces cerevisiae)* as the chassis. The fusion of FPPS to three different terpene synthases was examined and compared to the corresponding ‘free’ enzyme controls. The intracellular level of each enzyme was quantified using whole-cell proteomics, an approach that has not been harnessed in previous studies on enzyme fusion. We found the impact to be very enzyme-dependent: significant titre improvements were realised for three different isoprenoid pathways, but at varying magnitudes. Whole-cell proteomics analyses further revealed that – at least in the case of terpene synthases – increased *in vivo* enzyme stability from protein fusion is likely to be a greater contributor to the catalytic improvement than enzyme proximity.

## METHODS

### Molecular cloning and strain generation

All restriction enzymes and cloning reagents were purchased from New England Biolabs (NEB) unless specified otherwise. Cloning was performed using the isothermal assembly method (NEBuilder HiFi DNA Assembly Master Mix, NEB #E2621). The vector was prepared by excising the GFP-MIOX gene fragment from a P_GAL10_-GFP-MIOX^22^ yeast integrative plasmid by restriction enzyme digestion (ClaI + XhoI) followed by gel purification. Fragments with the appropriate 5’ and 3’ overhangs were generated by PCR and co-assembled with the linearised vector (refer to Supporting Information for PCR primers, templates, and synthetic gene sequences). The FPPS + nerolidol synthase (NES) and FPPS-NES constructs were built first, and the rest of the constructs generated by replacing either the FPPS or NES genes. The FPPS and NES regions were excised by double digesting with BglII + BamHI and SacI + XhoI, respectively. For the FPPS + deadFPPS-NES construct, the FPPS + NES plasmid was linearised at the SacI site and deadFPPS was inserted using a dsDNA gene block (synthesized by Integrated DNA Technologies Inc.). Isothermal assembly was performed by incubating at 50 °C for 1 hour and transformation into chemically competent *E. coli* DH5α (Zymo Mix & Go *E. coli* Transformation Kit, Zymo Research #T3001). Each construct was verified by Sanger sequencing (Australian Genome Research Facility) and next-generation sequencing (Addgene Inc.). All constructs were linearised by SwaI digestion prior to yeast transformation using the standard LiAc/SS carrier DNA/PEG method^23^. Yeast transformants were verified by colony PCR using cloning primers of the corresponding heterologous terpene synthase. Strains were cultured overnight in YPD medium (1% w/v yeast extract, 2% w/v peptone, 2% w/v glucose) and stored as 20% v/v glycerol stocks at −80 °C. At least 3 individual transformed colonies were maintained as biological replicates, with the exception of the FPPS + deadFPPS-NES construct where only 2 transformants were successfully recovered; for this construct, one of the transformants was cultured in two separate flasks and treated as biological replicates. Detailed descriptions of the yeast strains used in this work are provided in the table below.

**Table 1.** Description of *S. cerevisiae* strains used in this work. Unless otherwise indicated, the linker peptide sequence for the fusion protein constructs is AGGGGTGGA. Where available, the Addgene accession code for the yeast integrative plasmid (YIp) used to construct the strain is provided.

| Strain name | Description | Genotype description | Reference |
| --- | --- | --- | --- |
| CEN.PK2-1C | Base strain | MATa ura3-52 leu2-3,112 trp1-289 his3Δ1 MAL2-8c SUC2 | Entian and Kötter 2007<sup>24</sup> |
| o57BR | Base strain for terpene production with upregulated mevalonate pathway | CEN.PK2-1C derivative;<br>HMG2<sup>K6R</sup>(-152,-1)::HIS3-T<sub>EFM1</sub>-Ef.mvaS-P<sub>GAL1</sub>-P<sub>GAL10</sub>-ACS2-T<sub>ACS2</sub>-P<sub>GAL2</sub>-Ef.mvaE-T<sub>EBS1</sub>-P<sub>GAL7</sub>;<br>pdc5(-31,94)::P<sub>GAL2</sub>-ERG12-T<sub>NAT5</sub>-P<sub>TEF2</sub>-ERG8-T<sub>IDP1</sub>-T<sub>PRM9</sub>-MVD1-P<sub>ADH2</sub>-T<sub>RPL15A</sub>-IDI1-P<sub>TEF1</sub>-TRP1;<br>ERG9(1333, 1335)::T<sub>URA3</sub>-P<sub>GAL7</sub>-MVD1-T<sub>PRM9</sub>-P<sub>GAL2</sub>-ERG12-T<sub>NAT5</sub>-T<sub>IDP1</sub>-ERG8-P<sub>GAL10</sub>-P<sub>GAL1</sub>-IDI1-T<sub>RPL15A</sub>-P<sub>ACS2</sub>-SKP1-Os.TIR1; | Hayat et al. 2021<sup>25</sup> |
|  |  | MIG1(-1,1)::CUP1-AID* |  |
| Empty vector | Base strain transformed with an empty cassette containing only a *K. lactis* URA3 selection marker. | o57BR derivative;<br>ura3(1,704)::KIURA3 | This work;<br>plasmid pILGH4 from Peng et al. 2015<sup>26</sup> |
| FPPS + NES | Expresses FPPS (ERG20) and nerolidol synthase as separate, 'free' proteins | o57BR derivative;<br>ura3(1,704)::KIURA3-T<sub>HIS5</sub>-NES-P<sub>GAL7</sub>-T<sub>GAL10</sub>-FPPS- P<sub>GAL10</sub> | This work<br>Addgene #182482 |
| FPPS-NES | Expresses FPPS fused to nerolidol synthase, separated by a short linker peptide. | o57BR derivative;<br>ura3(1,704)::KIURA3-T<sub>HIS5</sub>-(FPPS-NES)-P<sub>GAL10</sub> | This work<br>Addgene #182485 |
| FPPS + deadFPPS-NES | Expresses FPPS and an inactive FPPS [ERG20(K197G-K254A)<sup>27</sup>] fused to nerolidol synthase. | o57BR derivative;<br>ura3(1,704)::KIURA3-T<sub>HIS5</sub>-[FPPS(K197G-K254A)-NES]-P<sub>GAL7</sub>-T<sub>GAL10</sub>-FPPS- P<sub>GAL10</sub> | This work<br>Addgene #182492 |
| FPPS + GFP-NES | Expresses FPPS and green fluorescent protein fused to nerolidol synthase. | o57BR derivative;<br>ura3(1,704)::KIURA3-T<sub>HIS5</sub>-(GFP-NES)-P<sub>GAL7</sub>-T<sub>GAL10</sub>-FPPS- P<sub>GAL10</sub> | This work<br>Addgene #182491 |
| FPPS + PTS | Expresses FPPS and patchoulol synthase as separate proteins | o57BR derivative;<br>ura3(1,704)::KIURA3-T<sub>HIS5</sub>-PTS-P<sub>GAL7</sub>-T<sub>GAL10</sub>-FPPS- P<sub>GAL10</sub> | This work<br>Addgene #182483 |
| FPPS-PTS | Expresses FPPS fused to patchoulol synthase, separated by a short linker peptide. | o57BR derivative;<br>ura3(1,704)::KIURA3-T<sub>HIS5</sub>-(FPPS-PTS)-P<sub>GAL10</sub> | This work<br>Addgene #182486 |
| FPPS + LS | Expresses FPPS and limonene synthase as separate proteins | o57BR derivative;<br>ura3(1,704)::KIURA3-T<sub>HIS5</sub>-LS-P<sub>GAL7</sub>-T<sub>GAL10</sub>-FPPS- P<sub>GAL10</sub> | This work |
| FPPS-LS | Expresses FPPS fused to limonene synthase, separated by a short linker peptide. | o57BR derivative;<br>ura3(1,704)::KIURA3-T<sub>HIS5</sub>-(FPPS-LS)-P<sub>GAL10</sub> | This work |
| FPPS(F96W-N127W) + LS | Expresses FPPS (with the F96W-N127W mutation which increases GPP release<sup>8</sup>) and | o57BR derivative; | This work<br>Addgene #182484 |
|  | nerolidol synthase as separate proteins | ura3(1,704)::KIURA3-T<sub>HIS5</sub>-LS-P<sub>GAL7</sub>-T<sub>GAL10</sub>-FPPS(F96W-N127W)-P<sub>GAL10</sub> |  |
| FPPS(F96W-N127W)-LS | Expresses FPPS (with the F96W-N127W mutation which increases GPP release<sup>8</sup>) fused to limonene synthase, separated by a short linker peptide. | o57BR derivative;<br>ura3(1,704)::KIURA3-T<sub>HIS5</sub>-[FPPS(F96W-N127W)-LS]-P<sub>GAL10</sub> | This work<br>Addgene<br>#182487 |

### Two-phase flask fermentation

Dodecane was used as an inert organic overlay in liquid culture to trap volatile terpenoids and minimise product evaporation^28^. Glycerol stocks stored at −80 °C were recovered on uracil drop-out agar plates. Cells were pre-cultured overnight in YPD medium. Pre-cultures were used to inoculate 20 ml YPDG medium (1% w/v yeast extract, 2% w/v peptone, 2% w/v galactose, 0.5% w/v glucose) at OD_600_ = 0.05 in 50 ml unbaffled shake flasks. 2 ml sterile-filtered dodecane was added before covering the flasks with foil. Cultures were incubated at 30 °C, 200 rpm shaking for 120 hours. Cultures were sampled periodically for OD_600_ measurement. Dry cell mass values were estimated by multiplying the OD_600_ reading at 72 h by a published conversion factor (0.644 g DCM/L per unit OD_600_)^29^. At 72 hours post-inoculation (h p.i.), samples of the dodecane layer and cell culture were taken for metabolite analysis and whole-cell proteomics. Cells were first pelleted and washed once with distilled water before storage. All dodecane and cell samples were stored at −20 °C until further analysis.

### High-performance liquid chromatography (HPLC)

Nerolidol, linalool, farnesol, geraniol, and geranylgeraniol were quantified simultaneously by HPLC as previously reported^30^, with minor changes. Briefly, dodecane overlay samples were thawed completely and then diluted 1:40 in ethanol (5 μl sample + 200 μl ethanol) in glass vials. From each sample, 20 μl was injected through a Waters Sunfire C18 column (3.5um, 4.6 × 150 mm, PN: 186002554) with a guard column (SecurityGuard Gemini C18, PN: AJO-7597). Separations were run either on an Agilent 1200 HPLC system or Thermo Fisher Vanquish HPLC system. Analytes were eluted at 0.9 ml/min at 35 °C. Solvent A was high-purity water, and solvent B was 45% acetonitrile + 45% methanol + 10% water. Elution was with a linear gradient of 5–100% solvent B from 0 to 24 min, 100% from 24 to 30 min, and 5% from 30.1 to 35 min. Analytes of interest were monitored using a diode array detector (Agilent DAD SL, G1315C) at 202 nm wavelength. The peak retention times are as follows: geraniol 21.4 min, linalool 21.6 min, nerolidol 26.6 min, farnesol 26.4 min, and geranylgeraniol 31.0 min. Sample concentrations were calculated using Chromeleon 7 (Thermo Scientific) by fitting the peak areas to a standard dilution curve. Each peak was manually inspected, and the peak region adjusted as required. Concentration data was exported into a Microsoft Excel spreadsheet for further analysis.

### Gas chromatography-mass spectrometry (GC-MS)

Patchoulol was analysed by GC-MS/MS on a Shimadzu GC/MS-TQ8050 NX system. The patchoulol standard was purchased in lyophilised form from Chengdu Alfa Biotechnology Co. Ltd. (patchouli alcohol >98%, #AB0523-0020) and dissolved in absolute ethanol. A sample of 1 μl (dodecane overlay) was injected into the GC inlet set at 250 °C in split mode of 1:5. Chromatographic separation was achieved using a Shimadzu Rxi-624Sil MS capillary column (20 m × 0.18 mm × 1 μm). Oven conditions were set at 100 °C starting temperature, then ramped at 20 °C/min to 270 °C and held for 1 min. Helium was used as the carrier gas at a flow rate of 0.6 mL/min. Compounds were fragmented by electron impact ionization and analysed in selected ion monitoring (SIM) mode, monitoring for patchoulol fragments with masses 138, 222, 161, and 179 m/z. Concentration data was exported into a Microsoft Excel spreadsheet for further analysis.

### Whole-cell proteomics by liquid chromatography-mass spectrometry (LC-MS)

Frozen cell pellets (from 2 ml culture collected at 72 h p.i.) were thawed on ice and resuspended with 100 μl phosphate buffered saline (PBS). Further processing was performed using 20 μl of resuspended sample. Trypsin digestion and sample clean-up was performed using S-Trap Micro Spin columns (ProtiFi) according to the manufacturer’s protocol, with minor changes. Samples were solubilised in 50 μl S-Trap lysis buffer (5% sodium dodecyl sulphate (SDS) in 50 mM Tris), before being reduced by adding 4 μl of 500 mM of dithiothreitol (DTT) and heating at 70 °C for 60 min. To alkylate cysteine residues, 8 μl of 500 mM iodoacetamide was added and samples were incubated in the dark for 30 min. Phosphoric acid (2.5 μl) was added (final concentration 12%), followed by 165 μl of S-Trap binding buffer (90% methanol in 100 mM Tris). The sample mix was then centrifuged through the S-Trap column at 4,000 g for 1 min. Three washes with 150 μl S-Trap binding buffer were performed, with centrifugation each time. Peptide digestion was initiated by adding 25 μl of 50 mM ammonium bicarbonate buffer (pH 8) containing 2 μg trypsin (Sequencing Grade Modified Trypsin, Promega #V5117) directly on top of the column and incubating overnight at 37 °C in a drying oven. Peptides were eluted by centrifuging successively with 40 μl of 5%, 50%, and 75% acetonitrile (all in 0.1% formic acid). Eluted peptides were dried down using a vacuum concentrator (Concentrator Plus, Eppendorf) and resuspended in 5% acetonitrile in 0.1 % formic acid prior to liquid chromatography.

The peptides were separated using an Ultimate 3000 RSLCnano HPLC system (Thermo Fisher Scientific, Germany) coupled to a Thermo Orbitrap Q Exactive HF Hybrid Quadrupole-Orbitrap Mass Spectrometer (Thermo Fisher Scientific, USA). For each sample, 2 μl was injected through a nanoEase™ M/Z CSH C18 column (130Å, 1.7μm, 300 μm × 100 mm) (Waters Corporation) via a desalting trap column at a flow rate of 20 μl/min. Peptides were eluted using a 60 mins gradient of 8% to 95% acetonitrile in 0.1% formic acid at 3 μl/min flow rate. The MS parameters were as follows: resolution = 60,000 and scan range = 100-800 m/z for MS1; isolation window = 2.0 m/z, TopN = 40, resolution = 75,000, nAGC target = 1e6 and max IT = 40 ms for MS2. Post-analysis was performed using Proteome Discoverer v2.4 (Thermo Fisher Scientific). For each experiment, a custom protein database was constructed by appending the sequences of heterologous proteins to a *S. cerevisiae* reference database (strain CEN.PK113-7D). Protein abundance data was exported into a Microsoft Excel spreadsheet for further analysis.

### Data analysis and plotting

Data collation and concentration upgrade calculations were performed with Microsoft Excel. Charts were plotted using the Matplotlib package in Python 3.

## RESULTS AND DISCUSSION

Isoprenoid biosynthesis in yeast starts with the mevalonate pathway, which produces the universal isoprenoid precursors isopentyl diphosphate (IPP; C_5_) and dimethylallyl diphosphate (DMAPP; C_5_)^31^ (Figure 1a). Successive condensation of IPP and DMAPP generate the prenyl diphosphate intermediates geranyl diphosphate (GPP; C_10_), farnesyl diphosphate (FPP; C_15_), and geranylgeranyl diphosphate (GGPP; C_20_), which are converted into diverse isoprenoid products by a class of enzymes called terpene synthases. In this work, we employed a base strain with a highly upregulated mevalonate pathway (o57BR)^25^ to provide a large supply of IPP and DMAPP for isoprenoid bioproduction.

**Figure 1.**
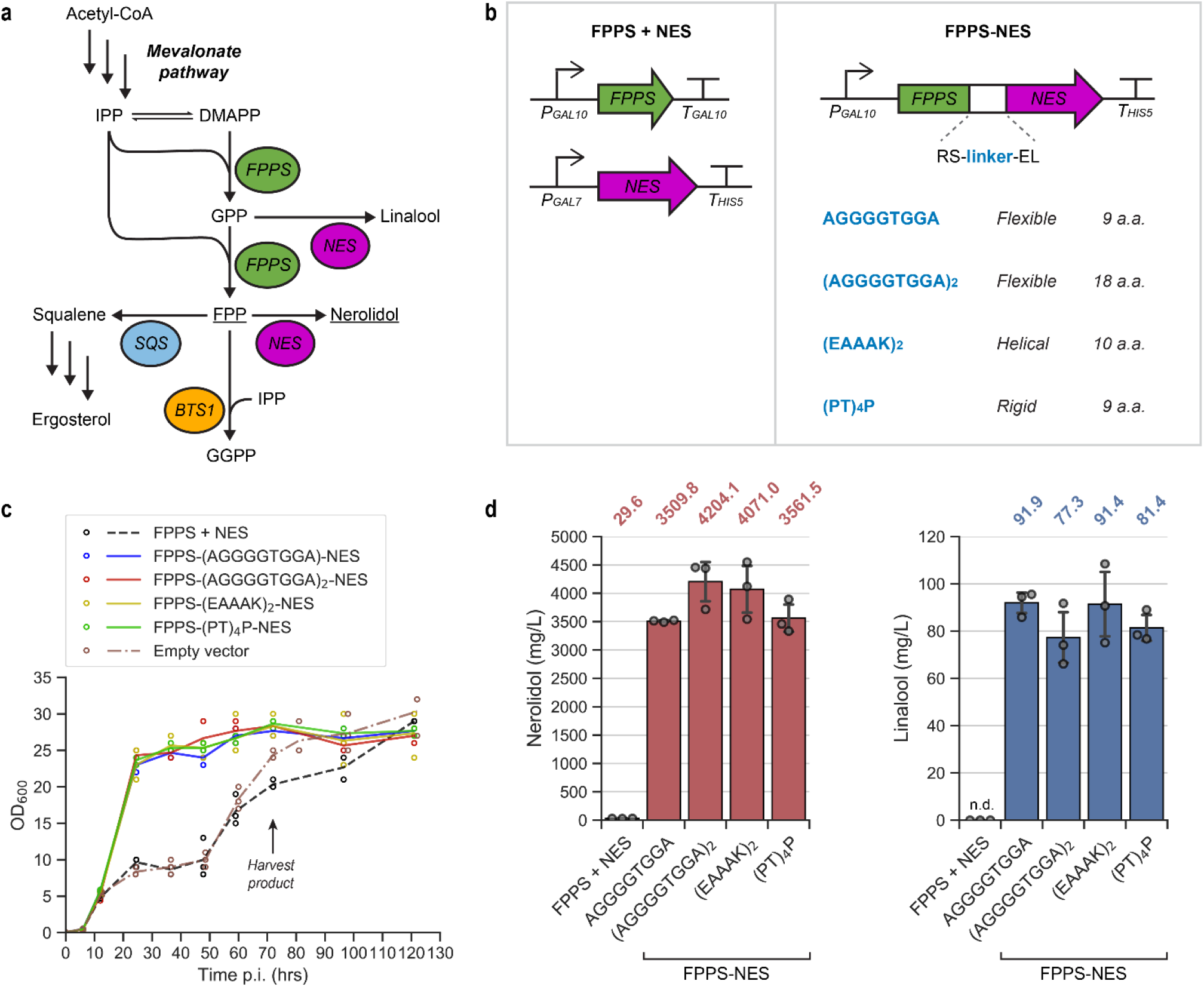
Fusion of farnesyl diphosphate synthase (FPPS) with nerolidol synthase (NES). **(a)** The base strain contains a highly upregulated mevalonate pathway^25^, which produces the universal isoprenoid precursors isopentyl diphosphate (IPP) and dimethylallyl diphosphate (DMAPP). Native yeast FPPS produces the isoprenoid intermediates geranyl diphosphate, GPP (minor product), and farnesyl diphosphate, FPP (major product, underlined). Introduction of NES and an overexpressed copy of FPPS redirects metabolic flux towards the production of the major NES product nerolidol (underlined). This variant of NES is promiscuous and also produces linalool as a minor product. The endogenous enzymes squalene synthase (SQS) and geranylgeranyl diphosphate synthase (BTS1) also compete for FPP, converting it into squalene and geranylgeranyl diphosphate (GGPP), respectively. **(b)** Schematic showing the expression cassettes for the free enzyme control (FPPS + NES) and enzyme fusion (FPPS-NES) constructs. Four peptide linkers were tested for the fusion configuration, with the sequences shown in blue. The labelled length (number of amino acids, a.a.) excludes the four residues (N-terminal RS and C-terminal EL) flanking the linker that are contributed by restriction enzyme sites. **(c)** Growth curves (OD_600_ values) over 120 hours post-inoculation (p.i.). The means of three biological replicates is plotted on the line graph with the individual data points shown as circles. **(d)** Nerolidol and linalool titres at 72 hours p.i., in mg product per L liquid culture. Bar chart values (also indicated above each bar) are means of three biological replicates, with error bars of 1 +/− STD. The individual data points are shown as circles, and ‘n.d.’ = not detected. Nerolidol titres normalised to dry cell mass (mg/g DCM) are provided in Figure S1.

### NES-FPPS fusion greatly improves nerolidol production and rescues cell growth

To generate strains that produce the sesquiterpene nerolidol, we introduced cassettes for galactose-inducible expression of yeast FPPS *(ERG20)* and nerolidol synthase (*NES*)^32^ from golden kiwifruit (*Actinidia chinensis*) into the genome. FPPS catalyses the two-step condensation of DMAPP with two IPP molecules to FPP *via* GPP (Figure 1a). Yeast FPPS releases a small proportion of GPP which can be utilised by heterologous enzymes^33^. The variant of NES we used is promiscuous and can also convert GPP into the side product linalool (at 68% efficiency of the FPP ➔ nerolidol reaction^32^). FPP is also consumed by the native enzymes squalene synthase (SQS/ERG9) and geranylgeranyl diphosphate synthase (GGPPS/BTS1)^31^.

Since peptide linker composition and length in enzyme fusions was previously reported to affect terpene production^4–6,21^, we designed the FPPS-NES fusion constructs with four different peptide linkers: flexible linkers AGGGGTGGA and (AGGGGTGGA)_2_, helical linker (EAAAK)_2_, and rigid linker (PT)_4_P (Figure 1b). The fusions were compared against a ‘free’ enzyme control that expresses FPPS and NES as separate proteins. Two different GAL promoters were used in the FPPS + NES cassette to avoid inadvertent homologous recombination during strain construction.

The control strains expressing the FPPS + NES or the empty vector (selection marker only) cassettes grew slowly and achieved low biomass accumulation in the early phase of fermentation (<48h p.i.) (Figure 1c). This is consistent with previous reports of growth impairment in strains with engineered isoprenoid pathways in both *S. cerevisiae*^28,30,34,35^ and *E. coli*^36,37^. Over-accumulation of prenyl diphosphate intermediates (IPP, DMAPP, and/or FPP) resulting in cellular toxicity has been implicated as the basis of this growth impairment^38,39^. Interestingly, the FPPS-NES fusion strains exhibited growth profiles that were similar to strains without mevalonate pathway upregulation^22^, achieving ~2.5-fold higher biomass accumulation compared to the controls in the first 48 hours (Figure 1c). This observation suggests that fusion of NES to FPPS provides a mechanism to mitigate the toxicity observed in flux-enhanced strains.

Coexpressing FPPS and NES (without fusion) resulted in a modest nerolidol titre of 29.6 mg/L culture at 72 h p.i. (Figure 1d). Fusion of FPPS to NES delivered a remarkable increase in nerolidol of >110-fold for each of the four fusion constructs, resulting in titres of between 3.51 – 4.20 g/L culture, even under non-optimised shake-flask conditions. This is an unusually large fold improvement from a single engineering step, especially given that previous studies investigating the fusion of yeast FPPS with terpene synthases only saw titre improvements of <4-fold^4,6–8^.

No linalool could be detected in the free enzyme control (Figure 1d). In contrast, significant linalool production (77.3 – 91.9 mg/L) was observed in all FPPS-NES fusion strains. Less linalool was produced compared to nerolidol (approximately an order of magnitude less) because of two reasons: the small free pool of GPP produced by the bifunctional yeast FPPS^7,33^, and the lower preference of this NES variant for GPP compared to FPP (*k_cat_/Km* 69 s^-1^ mM^-1^ for GPP *versus* 300 s^-1^ mM^-1^ for FPP)^32^. No significant differences were observed in nerolidol or linalool production for the different peptide linker sequences, suggesting that the observed effect is enzyme-specific rather than linker-specific.

We also detected the C15 isoprenoid pathway side product farnesol (10.1 – 33.6 mg/L) in all cultures (Figure S1). In the presence of excess prenyl diphosphates – especially when the mevalonate pathway is upregulated – prenyl alcohols are commonly produced by the action of nonspecific phosphorylases^7,18^. Since the levels of farnesol were two orders of magnitude less than nerolidol, conversion to prenyl alcohols was not a significant source of carbon loss. Additionally, accumulation of DMAPP, GPP, and FPP are known to inhibit the enzyme mevalonate kinase (ERG12)^40^, exerting negative feedback control of the yeast mevalonate pathway. Poor conversion of FPP into nerolidol – such as in the FPPS + NES strain – could have led to downregulation of the mevalonate pathway, and further reduction in FPP synthesis. This would explain the limited farnesol production by the FPPS + NES strain, despite the low metabolic flux diversion through the nerolidol pathway.

### Mitigation of toxicity and increased production is due to protein stabilisation and not co-localisation of the pathway enzymes

Aside from changing enzyme spatial organisation, fusion of FPPS to NES could impact protein translation and/or *in vivo* stability. To separately assess the effects of enzyme co-localisation and enzyme stability, we designed an expression experiment where free FPPS was coexpressed with either deadFPPS-NES or GFP-NES. The deadFPPS protein is a mutated FPPS (K197G-K254A) with no catalytic activity^27^, and GFP is yeast-enhanced green fluorescent protein^41^ (Figure 2a). If co-localisation of the FPPS and NES active sites were the sole reason for the activity enhancement in the FPPS-NES fusion strains, then we would not expect to see an enhancement when NES is fused to an inactive protein. We chose deadFPPS as the fusion partner to investigate if the enhancement previously shown by the FPPS-NES constructs (Figure 1d) could be attributed to stabilisation of NES, and GFP because it is known to be a stabilising tag^42^.

**Figure 2.**
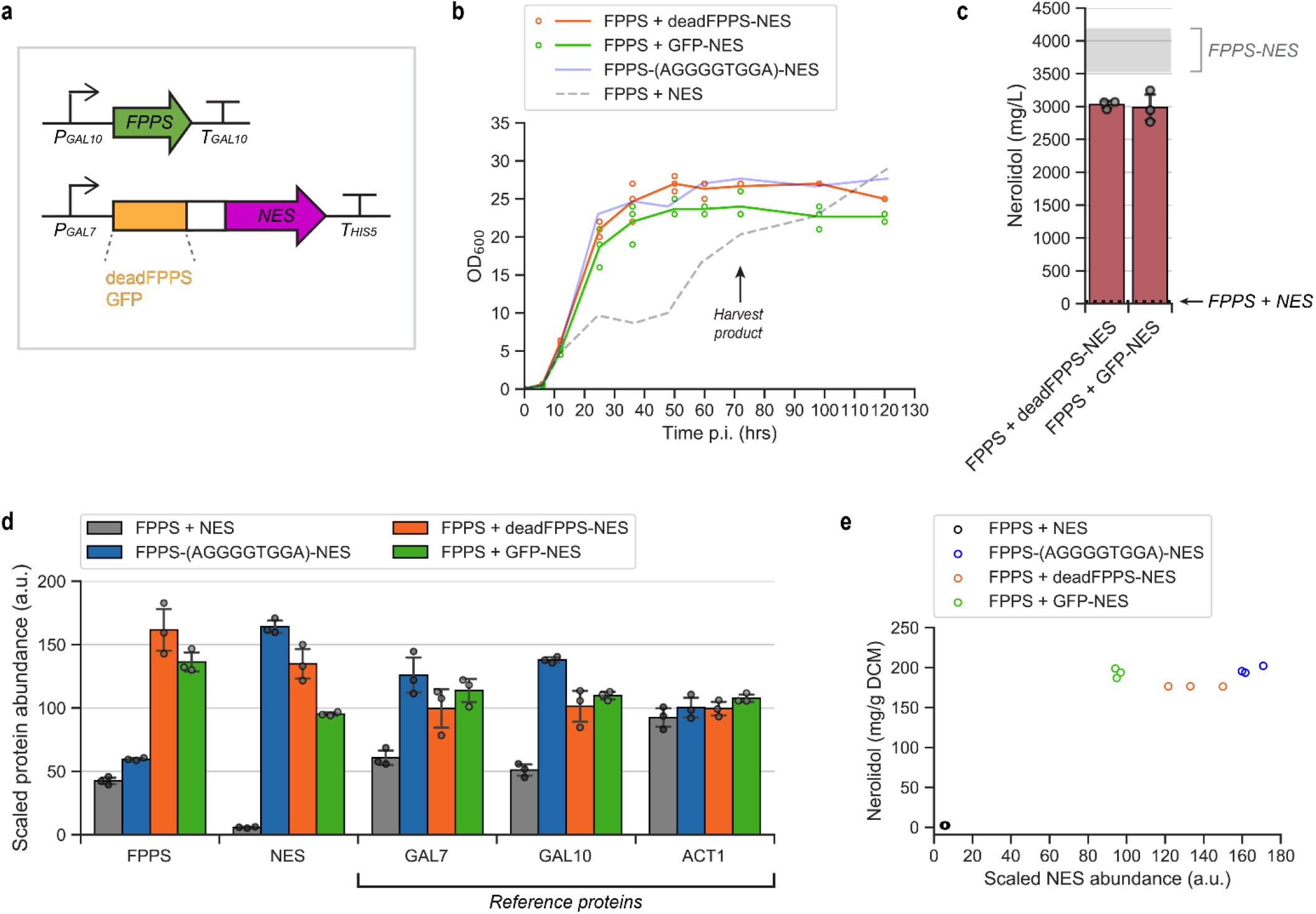
NES stabilisation by fusion to an inactive protein. **(a)** Expression cassette schematic for the coexpression of FPPS and NES fused either to an inactive FPPS mutant, deadFPPS (FPPS + deadFPPS-NES) or GFP (FPPS + GFP-NES). The AGGGGTGGA linker peptide was used to bridge NES and its fusion partner. **(b)** Growth curves (OD_600_ values) over 120 hours post-inoculation (p.i.). The means of three biological replicates are plotted on the line graph with the individual data points shown as circles. The growth curves of FPPS-(AGGGGTGGA)-NES and FPPS + NES (from Figure 1c) are overlayed for comparison. **(c)** Nerolidol titres at 72 h p.i., in mg product per L liquid culture. Values are means of three biological replicates with error bars of 1 +/− STD. The individual data points are shown as circles. The mean titre for FPPS + NES (black dotted line) and the range of mean titres across all FPPS-NES fusion strains (grey band) (from Figure 1d) are overlayed for comparison. Nerolidol titres normalised to dry cell mass (mg/g DCM) are provided in Figure S1. **(d)** Relative protein abundances of FPPS and NES as well as key reference proteins (native to the host cell) from whole-cell proteomics analysis, scaled to enable easier comparison across strains. Note that deadFPPS is indistinguishable from FPPS due to their high sequence identity. Values are means of three biological replicates with error bars of 1 +/– STD. **(e)** Nerolidol production per unit dry cell mass (mg/g DCM) versus scaled NES abundance.

Interestingly, the two new fusion strains had a growth pattern similar to the FPPS-NES fusion strains (Figure 2b). This suggests an alleviation of isoprenoid intermediate build-up even in the absence of enzyme co-localisation, thus disproving the hypothesis that co-localisation of sequential enzymes in this metabolic cascade is a key influence in the mitigation of toxicity. Both deadFPPS-NES and GFP-NES fusions produced high nerolidol titres of 3.03 g/L and 2.99 g/L, respectively (Figure 2c). These titres represent a >101-fold increase over free FPPS + NES expression, and approach those of the FPPS-NES fusions.

Additionally, these two constructs produced significant amounts of the longer chain prenyl alcohol geranylgeraniol (Figure S1), while levels of this side product were below the limit of quantification in FPPS + NES and FPPS-NES. Presuming that the detected geranylgeraniol came from nonspecific hydrolysis of excess GGPP^8,43^ (which in turn comes from BTS1 conversion of FPP; Figure 1a), this implies inefficient consumption of the FPP pool by NES compared to the FPPS-NES fusions. Farnesol was detected at levels comparable to that of FPPS-NES fusion strains (Figure S1).

To quantify protein expression, samples collected at 72 h p.i. were analysed using whole-cell proteomics (LC-MS/MS). NES abundance in all fusion constructs were massively elevated relative to the FPPS + NES control strain, specifically by 28-fold for FPPS-NES (AGGGGTGGA linker), 23-fold for deadFPPS-NES, and 17-fold for GFP-NES (Figure 2d; Table S3). There is a clear relationship between NES expression and nerolidol production (Figure 2e), suggesting that nerolidol production is mostly limited by NES levels. Our results also show that *A. chinensis* NES expression can be improved by fusion to a more stable partner. The high sequence identity between deadFPPS and wtFPPS (differing only in one of the detected peptides) makes them impossible to differentiate by LC-MS/MS; this explains the much higher apparent levels of FPPS in the FPPS + deadFPPS-NES strain compared to the FPPS-NES strain. However, we are unable to explain why detected FPPS appears to be similarly elevated in the GFP-NES strain.

The levels of native and engineered galactose-inducible proteins (expressed using GAL promoters) were found to be higher overall in the FPPS-NES fusion compared to the other NES configurations (Figure 2d; Figure S2), suggesting earlier switching-on of protein expression. The cell growth rate and the copy number of GAL promoters are two factors that could influence the timing of galactose induction. GAL promoters respond to the ratio of galactose and glucose^44^, and a slow-growing strain would take longer to deplete glucose in the culture medium (present at 0.5% w/v), leading to delayed gene expression. This is particularly evident in the slow-growing FPPS + NES strain, which consistently exhibits lower expression of galactose-inducible genes (Figure 2d; Figure S2). Additionally, all GAL promoters in yeast are regulated by the Gal4p-activator/Gal80p-repressor transcription factor complex^28,45^; cassettes engineered with multiple GAL promoters would thus share the same pool of transcription factors. This is a plausible explanation for the apparent slight delay in GAL induction in the deadFPPS-NES and GFP-NES constructs (which contain two GAL promoters) compared to the FPPS-NES fusion (which contains a single GAL promoter). These findings may have important implications for the use of GAL promoters for metabolic engineering and warrant further investigation.

Notably, the deadFPPS-NES and GFP-NES strains did not produce any detectable linalool. This suggests that enzyme co-localisation is a key driver for production of linalool, but not for nerolidol. Since FPPS can access GPP directly from its own active site, a strong ‘pull’ is required to divert GPP flux into any other pathway. High levels of NES did not appear to provide sufficient flux pull without co-localisation of FPPS and NES by enzyme fusion. In contrast, the native enzymes BTS1 and SQS do not appear to be very competitive with NES for FPP, as evidenced by the secretion of prenyl alcohols (which is a symptom of prenyl diphosphate accumulation^7,18^; Figure S1). Nevertheless, we cannot rule out any possible influences of deadFPPS or GFP fusion on the substrate specificity of NES.

Another instance where enzyme fusion affected the products of a promiscuous terpene synthase differentially has been reported in the literature. When GPPS and pinene synthase were expressed either as free enzymes or as a translationally fused protein in *E. coli*, a change in the ratio of isoprenoid products (α-pinene and β-pinene) was observed, an effect which the authors attributed to differences in GPP concentration^16^. GPPS from *Vitis vinifera* is known to be inhibited by a high concentration of GPP^46^, although it is unknown if the same is true for the yeast FPPS.

### Stabilisation of diverse terpene synthases by translational fusion

We next investigated if fusion to FPPS leads to a similar stabilisation of other terpene synthases to evaluate if our findings were specific to NES and/or the strain background that we used. We replaced NES in the free enzyme (FPPS + NES) and FPPS-(AGGGGTGGA)-NES cassettes with *Pogostemon cablin* patchoulol synthase (PTS)^47^ for the production of the sesquiterpene patchoulol (Figure 3a) and *Citrus limon* limonene synthase (LS)^35^ for the production of the monoterpene limonene (Figure 3d). In the integrative cassettes of the LS-expressing strains, wtFPPS was also replaced with a GPP-overproducing FPPS mutant [FPPS(F96W-N127W)^7^] to provide enough substrate for detectable limonene titres^48^.

**Figure 3.**
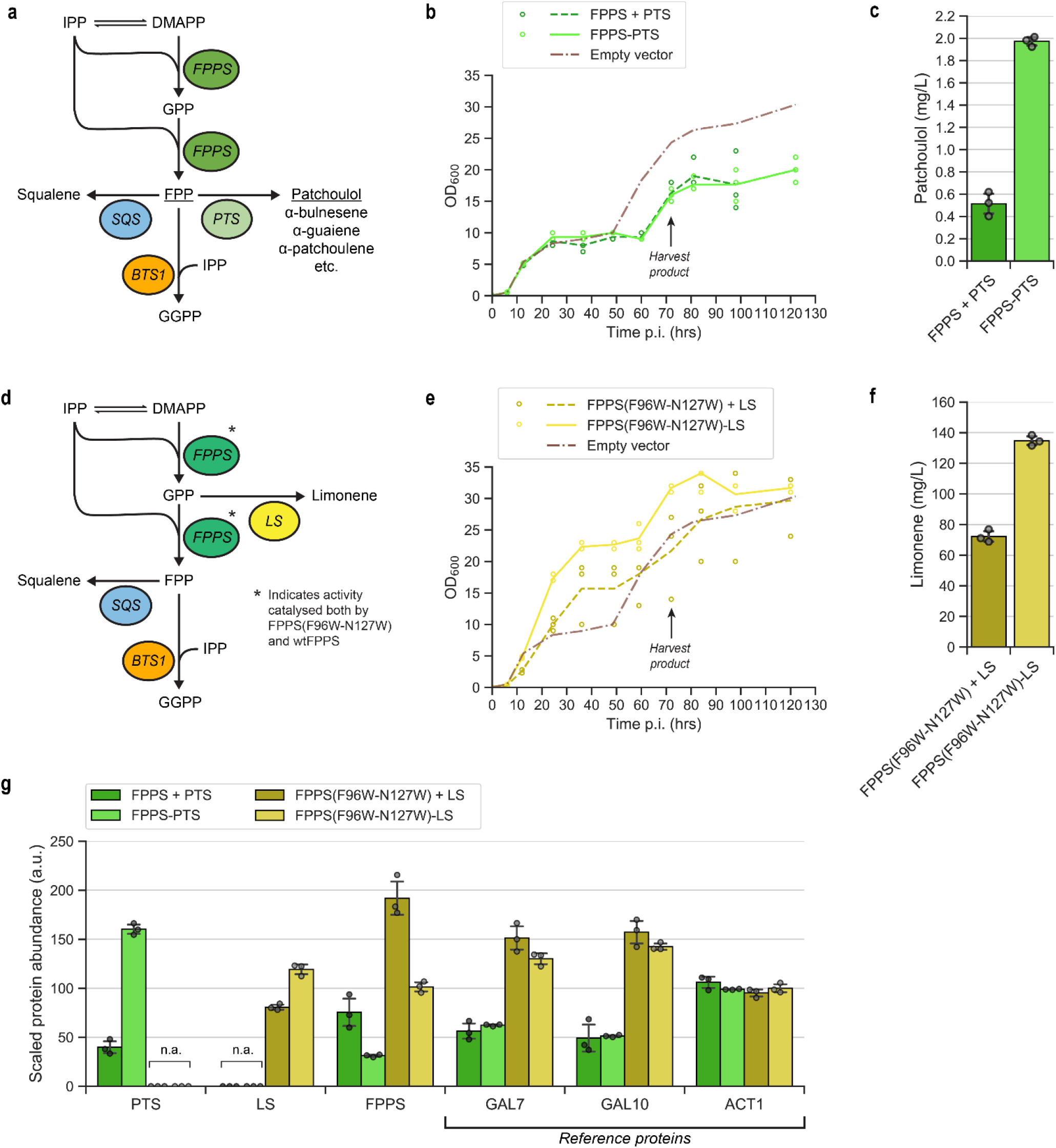
Fusions of FPPS with patchoulol synthase (PTS) and limonene synthase (LS). Reaction schematic showing the terpenoid production pathway for strains expressing **(a)** PTS or **(d)** LS. Growth curves (OD_600_ values) are shown over 120 hours post-inoculation (p.i.) for **(b)** PTS strains and **(e)** LS strains. The means of three biological replicates are plotted on the line graph with the individual data points shown as circles. The growth curve of the “Empty” vector strain (from Figure 1c) is overlayed for comparison. **(c)** Patchoulol and **(f)** limonene titres at 72 hours p.i., in mg product per L liquid culture. Values are means of three biological replicates with error bars of 1 +/−STD. The individual data points are shown as circles. The mean titre for FPPS + NES (black dotted line) and the range of titres across all FPPS-NES fusion strains (grey band) (from Figure 1d) are overlayed for comparison. **(g)** Relative protein abundances of PTS, LS and FPPS as well as key reference proteins from whole-cell proteomics analysis. Note that FPPS(F96W-N127W) is indistinguishable from FPPS due to their high sequence identity. Values are means of three biological replicates with error bars of 1 +/−STD.

For PTS and LS, enzyme fusion led to considerably less prominent improvements in growth and product titres compared to NES, indicating that the effect of enzyme fusion is enzyme-dependent. For PTS, both free enzyme (FPPS + PTS) and fusion (FPPS-PTS) strains exhibited similar, slow growth reminiscent of the FPPS + NES strain (Figure 3b). The reported turnover number (*k_cat_* for FPP) for *P. cablin* PTS is 0.43×10^-3^ s^-1^,^47^ which is three orders of magnitude slower than that of *A. chinensis* NES (*k_cat_* = 0.24 s^-1^)^32^. The slow kinetics of PTS suggests that PTS levels could be limiting in both the constructs we evaluated, creating an FPP flux bottleneck. In the case of LS, FPPS(F96W-N127W)-LS had slightly improved growth compared to the non-fusion control (Figure 3e).

No difference in growth between free and fused enzymes was seen for LS strains that overexpress wtFPPS instead of FPPS(F96W-N127W) (Figure S3), indicating that the growth effects are linked to the utilisation of prenyl diphosphate intermediates. This experiment also implicates FPP toxicity because strains overexpressing wtFPPS were more impaired in growth compared to the FPPS(F96W-N127W) strains (Figure 3e, Figure S3). Although limonene is known to be toxic to yeast, the levels observed here are far below the level required for growth inhibition^49^.

Enzyme fusion increased patchoulol production by 3.8-fold (Figure 3c), concomitant with a similar 4.0-fold increase in intracellular PTS (Figure 3g, Table S3). Terpene synthase expression also scaled closely with isoprenoid production in the case of LS, where enzyme fusion increased LS levels by 1.5-fold (Figure 3g) and limonene titres by 1.9-fold (Figure 3f, Table S3). However, it is unclear why fusion to LS or PTS led to a decrease in FPPS expression (Figure 3f), while the opposite was true in the case of NES (Figure 2d). Yeast FPPS could be sensitive to translational fusion at the C-terminal, as also shown by a recent study (in the green alga *Chlamydomonas reinhardtii*), where higher bisabolene production was observed when a fluorescent protein was fused to the N-terminal of yeast FPPS compared to when the fluorescent protein was fused to the FPPS C-terminal^50^.

The observation that translational protein fusion can improve terpene synthase expression has been reported in several other works. Limonene production was increased 40% in Chinese baijiu fermentation when LS was fused to neryl disphosphate synthase, with the length of the linker peptide and order of the fused enzymes strongly influencing titres^21^. In a cyanobacterial system (*Synechocystis* sp. PCC 6803), β-phellandrene synthase expression was greatly increased upon fusion with a native protein (cpcB), leading to a 100-fold improvement in β-phellandrene yield (3.2 mg/g DCM)^51^. However, it was unclear at that time if the effect could be extended to different terpene synthases as well as to other host organisms. The addition of C-terminal tags to a geraniol synthase-FPPS(F96W-N127W) fusion protein also increased protein expression and improved geraniol production in yeast^52^. In a *Chlamydomonas reinhardtii* study, levels of FPPS and SQS-like enzymes were elevated by modification of the N-end terminal residues, which suggested decreased protein degradation by the N-end rule^53^.

## CONCLUSIONS

Our work sheds light on the mechanism of titre enhancement from enzyme fusion, especially in the case of terpene synthases. The effect of enzyme fusion on enzyme stability has not previously been identified as a significant contributing factor, with previous works emphasising the assumed role of inter-enzyme proximity. This is a conspicuous gap in the literature, especially since translational fusion to other proteins and/or the modification of N-terminal residues is widely known to affect protein expression.

Here, we observed a strong relationship between intracellular terpene synthase levels and isoprenoid product titres. Although the terpene synthase-catalysed step is generally one of the limiting steps in isoprenoid bioproduction, this is often attributed to the slow catalytic rates of terpene synthases^28,54^. Our observations show that terpene synthase expression and/or stability can also limit productivity. This would explain why increasing the gene copy number of the corresponding terpene synthase is particularly effective for improving limonene and nerolidol production^48^. Furthermore, our work suggests that heterologously expressed terpene synthases should additionally be screened for poor expression or *in vivo* protein instability.

Another interesting observation is that significant linalool production was achieved only when FPPS was translationally fused to NES. Unlike the case of nerolidol, upregulation of NES expression alone did not measurably affect linalool titres. This suggests a potential impact of enzyme spatial organisation on the behaviour of promiscuous enzymes such as NES.

Stabilisation of terpene synthases by fusion to more stable domains has been identified as an excellent method of increasing terpene synthase levels. This approach can be used instead of, or in combination with, traditional metabolic engineering methods such as using a strong gene promoter or introducing additional gene copies.

## ASSOCIATED CONTENT

Supporting information file (PDF).

## ACKNOWLEDGEMENTS

L.C.C. and additional research costs for this work were supported by a CSIRO-UQ strategic fund under the Synthetic Biology Future Science Platform. F.S. and B.P. acknowledge support from CSIRO in the form of a Synthetic Biology Future Science Platform Fellowship. B.P. acknowledges support from the Australian Research Council Centre of Excellence in Synthetic Biology (project number CE200100029), which is funded by the Australian Government. Metabolomics Australia is supported by BioPlatforms Australia through the Commonwealth Government’s National Collaborative Research Infrastructure Strategy (NCRIS). The authors thank Mitchell J. O’Sullivan (Queensland University of Technology) for assistance with writing Python scripts for data analysis as well as Gert Talbo and Timothy McCubbin (Metabolomics Australia, Queensland Node) for assistance with the interpretation of whole-cell proteomics data. The authors are grateful to Colin Scott (CSIRO) for helpful discussions and experiment suggestions.

## AUTHOR CONTRIBUTIONS

L.C.C. and F.S designed experiments. L.C.C generated strains and conducted experiments. L.L. performed whole-cell proteomics analysis by LC-MS/MS. T.S. developed the method for and performed patchoulol analysis by GC-MS/MS. M.R.P. performed metabolite analysis by HPLC. B.P. developed the base strain for nerolidol production and assisted with experimental design and data interpretation. Z.L. assisted with experiments and HPLC data analysis. F.S., G.S., and C.E.V. supervised the project. L.C.C., C.E.V., F.S., and G.S. wrote the manuscript. All authors read and approved the final manuscript.

