## Supplementary Information for "Translational fusion of terpene synthases enhances metabolic flux by increasing protein stability"

**Supplementary Figures and Tables:**

*Figure S1. Levels of key isoprenoid pathway side products at 72 h fermentation (HPLC).* 3

*Figure S2. Whole-cell proteomics of NES strains (LC-MS/MS).* 4

*Figure S3. Growth curves of strains expressing LS and wtFPPS.* 5

*Figure S4. Whole-cell proteomics of PTS and LS strains (LC-MS/MS).* 6

*Table S1. PCR primer sequences for strain construction.* 7

*Table S2. Synthetic gene sequences.* 11

*Table S3. Comparing the relative levels of terpene synthase and the corresponding isoprenoid product.* 16

**Figure S1.** Levels of key isoprenoid pathway side products at 72 h fermentation (HPLC).

Titres are presented in mg product per L cell culture. Values are means of 3 biological replicates with error bars of 1 +/- STD. The individual data points are shown as circles.

(**a**) Farnesol levels of NES-expressing strains. (**b**) Geranylgeraniol levels of strains coexpressing FPPS and deadFPPS-NES / GFP-NES. For the free enzyme control (FPPS + NES) and FPPS-NES fusion strains, geranylgeraniol was below the limit of quantification and therefore not shown. (**c**) Nerolidol titres of NES-expressing strains, presented per unit dry cell mass (mg/g DCM). Dry cell mass values were estimated by multiplying the $OD_{600}$ reading at 72 h by a published conversion factor (0.644 g DCM/L per unit $OD_{600}$; ref. Myers et al. 2013[1]). The mean titre for FPPS + NES is 2.26 mg/g DCM. (**d**) Farnesol levels of LS-expressing strains. The two 'FPPS' strains on the right are strains expressing wtFPPS (refer to Figure S3 for growth curves). (**e**) Geraniol levels of LS-expressing strains. No geraniol was detected in the strains expressing wtFPPS.

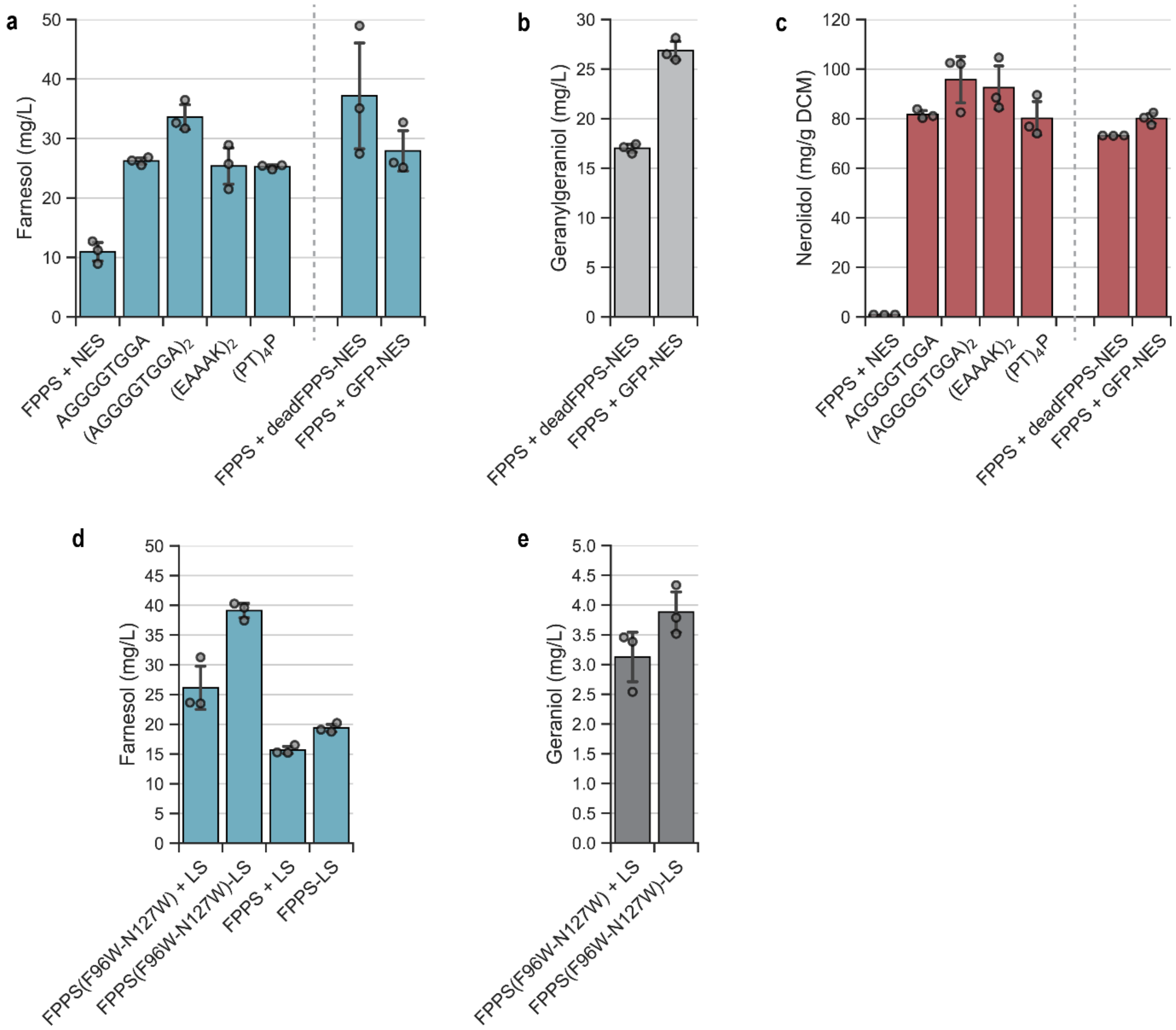

**Figure S2.** Whole-cell proteomics of NES strains (LC-MS/MS).

'Normalised protein abundance' is a measure of the relative detected levels of each protein in the sample, normalised to the total protein for that sample (in arbitrary units). In 'Scaled protein abundance', the level of each protein is individually re-scaled so that the mean equals 100 arbitrary units across samples; this enables easier comparison especially for proteins detected at low abundance. Bar charts show the mean +/- 1 standard deviation, and the individual data points (3 biological replicates each). All proteins shown were detected with high confidence. (**a**) Engineered nerolidol pathway genes. For the FPPS + deadFPPS-NES construct, the peptides for FPPS and deadFPPS are indistinguishable except for one that was detected at relatively low abundance; this means that the abundance value for 'FPPS' includes both FPPS and deadFPPS. KlURA3 (*K. lactis* URA3) is the selection marker used in the NES cassette. EfmvaE, EfmvaS, and SKP1-OsTIR1 are heterologous genes in the base strain o57BR[2]. EfmvaE and EfmvaS were expressed using GAL2 and GAL1 promoters respectively, and thus can act as additional reporters for the degree of galactose induction in each strain. (**b**) The relative levels of native galactose-inducible proteins (GAL1, GAL2, GAL7, GAL10) could also be indicative of the stage of growth and degree of galactose induction in each strain. In the engineered nerolidol pathway, the GAL10 promoter was used for FPPS / FPPS-NES expression and GAL7 promoter for NES / deadFPPS / GFP-NES expression. (**c**) The relative levels of the housekeeping proteins ACT1, TEF1, and PGK1 were very similar except in FPPS + NES, which exhibited impaired growth. (**d**) Relative levels of un-engineered ergosterol pathway proteins. Only the proteins that were detected with high confidence and abundance are shown.

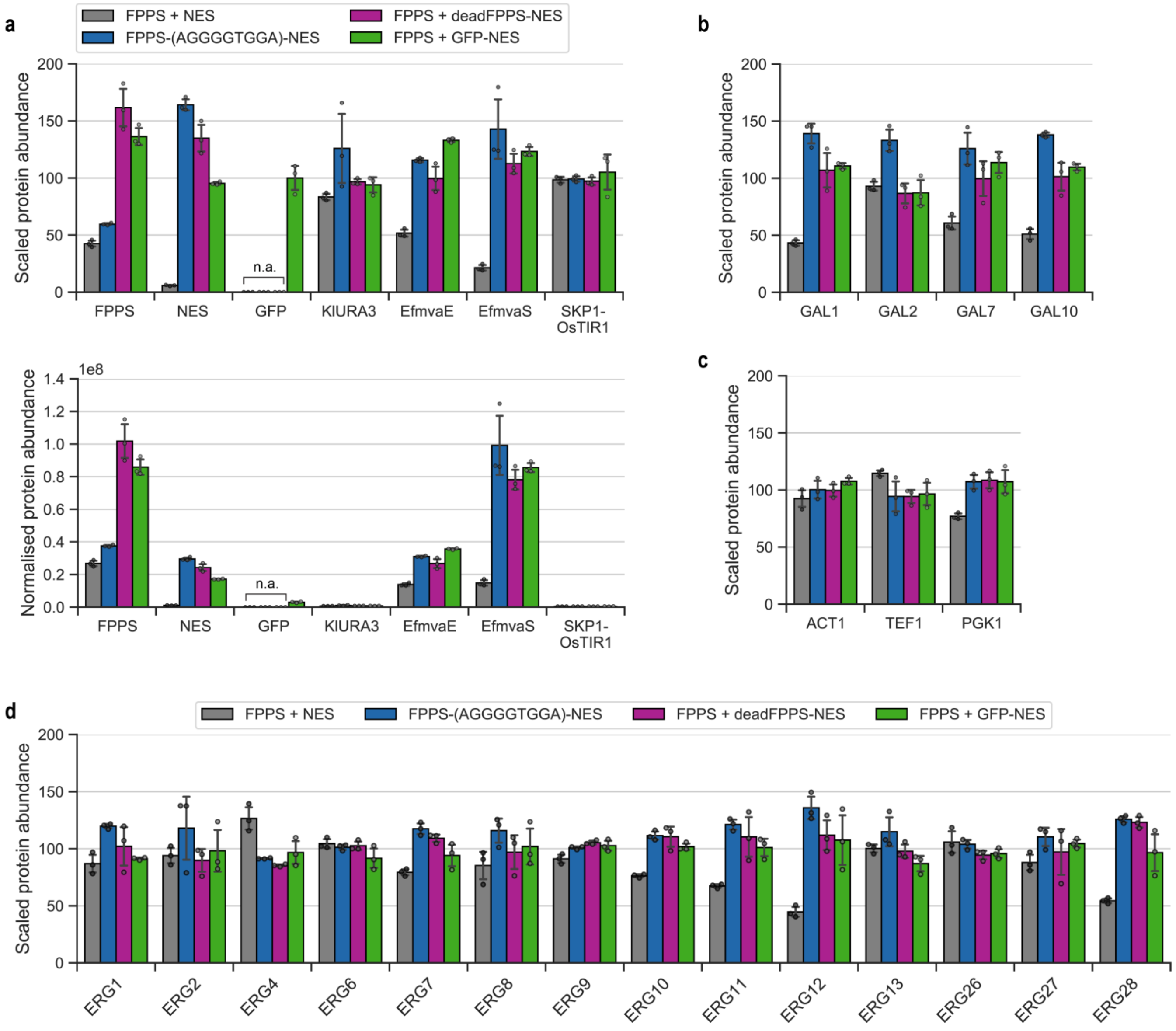

**Figure S3.** Growth curves of strains expressing LS and wtFPPS.

'Free enzyme' and enzyme fusion cassettes were designed to be identical to the limonene production cassettes used in Figure 3 in the main text, but with wild-type FPPS in place of FPPS(F96W-N127W). These strains did not produce any detectable limonene. Strains in this set displayed markedly slower growth compared to strains expressing FPPS(F96W-N127W), and the growth profiles are very similar to those of the PTS strains.

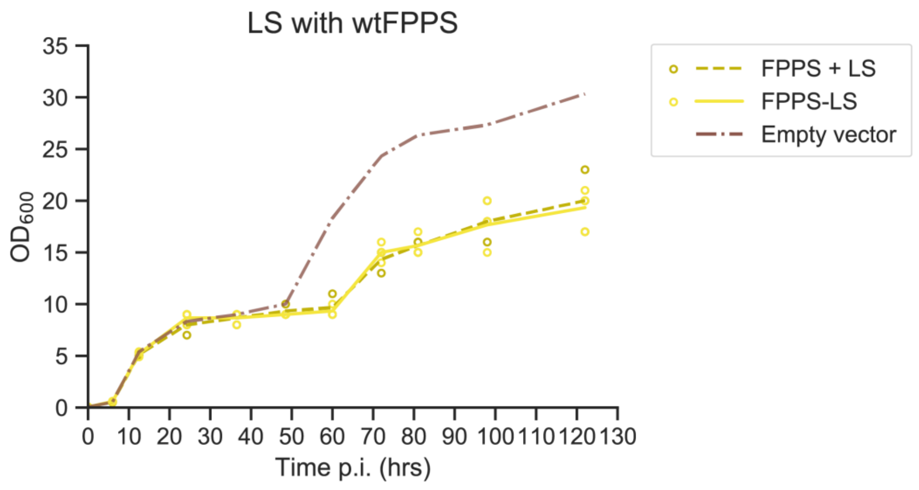

**Figure S4.** Whole-cell proteomics of PTS and LS strains (LC-MS/MS).

Bar charts show the mean +/- 1 standard deviation, and the individual data points (3 biological replicates each). All proteins shown were detected with high confidence. (**a**) Engineered terpene pathway genes. 'FPPS' includes both wtFPPS and FPPS(F96W-N127W). EfmvaE, EfmvaS, and SKP1-OsTIR1 were introduced in previous engineering of the base strain (o57BR). (**b**) The relative levels of native galactose-inducible proteins (GAL1, GAL2, GAL7, GAL10). In the engineered terpene production pathway, the GAL10 promoter was used for FPPS / FPPS-PTS / FPPS(F96W-N127W)-LS expression and GAL7 promoter for PTS / LS expression. (**c**) The relative levels of the housekeeping proteins ACT1, TEF1, and PGK1. (**d**) Relative levels of un-engineered ergosterol pathway proteins. Only the proteins that were detected with high confidence and abundance are shown.

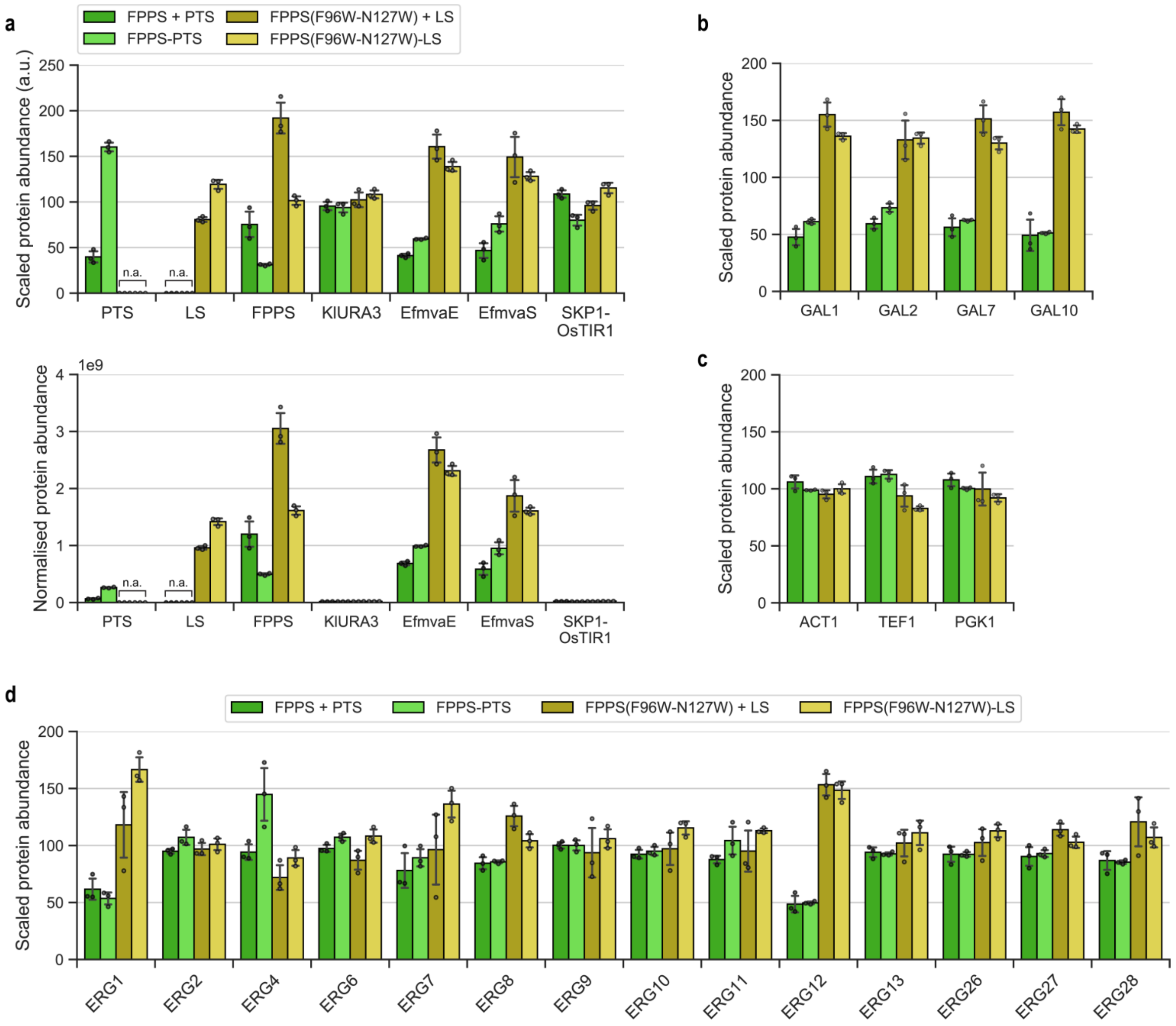

**Table S1.** PCR primer sequences for strain construction.

All gene constructs were cloned by isothermal assembly, which requires generating dsDNA fragments that overlap with adjacent fragments. Fragments with the appropriate 5' and 3' overlaps can be produced by mixing and matching PCR primers with suitable overhangs.

| Name | Description | Sequence (5' - 3') | Length (nt) | PCR template | PCR fragment generated | Constructs |
|---|---|---|---|---|---|---|
| ERG20_pGAL10_fw | FPPS forward, with overhang for $P_{GAL10}$ | AAAGTAAGAATTTTTGAAAATTCAATATAAGGATCCAAAATGGCTTCAGAAAAAGAAATT | 60 | pJT9R (Peng et al. 2017[3] ) for wtFPPS / pIT6EG7M (Peng et al. 2022[4]) for FPPS(F96W-N127W) | FPPS, FPPS(F96W-N127W) | All FPPS and FPPS(F96W-N127W) constructs |
| ERG20_tGAL10_rv | FPPS reverse, with overhang for $T_{GAL10}$ | CAAGAAGGATAGTAAGCTGGCAAAAGATCTTTATTTGCTTCTCTTGTAAACTTTGTTC | 58 | pJT9R for wtFPPS / pIT6EG7M for FPPS(F96W-N127W) | FPPS, FPPS(F96W-N127W) | FPPS + NES, FPPS(F96W-N127W) + LS |
| tGAL10_ERG20_fw | Forward primer for $T_{GAL10}$-$P_{GAL7}$ region, with overhang for FPPS | GAACAAAGTTTACAAGAGAAGCAAATAAAGATCTTTTGCCAGCTTACTATCCTTCTTG | 58 | *S. cerevisiae* S288C genomic DNA | $T_{GAL10}$-$P_{GAL7}$ region | FPPS + NES |
| pGAL7_AcNES1_rv | Reverse primer for $T_{GAL10}$-$P_{GAL7}$ region, with overhang for NES | CTGCTGCGGTAGCCATTTTGAGCTCTTTTGAGGGAATATTCAACTGTTTT | 50 | *S. cerevisiae* S288C genomic DNA | $T_{GAL10}$-$P_{GAL7}$ region | FPPS + NES |
| AcNES1_pGAL7_fw | NES forward, with overhang for $P_{GAL7}$ | GATAAAAAAAAACAGTTGAATATTCCCTCAAAAGAGCTCAAAATGGCTACCGC | 60 | pJT9R | NES | FPPS + NES |

| | | | | | | |
|---|---|---|---|---|---|---|
| | | AGCAGGT | | | | |
| AcNES1_HIS5t_rv | NES reverse, with overhang for $T_{HIS5}$ | AACTGTACATATACTGTTTAAATTAATCTATCTCGAGTTATAAAGATGTGTTATAGATCA | 60 | pJT9R | NES | All NES constructs |
| ERG20_AG4TGGA_rv | FPPS reverse for FPPS-NES fusion protein; appends linker peptide (AGGGGTGGA) | AGCACCACCAGTACCTCCTCCACCAGCAGATCTTTTGCTTCTCTTGTAAACTTTGTTC | 58 | pJT9R for wtFPPS / pIT6EG7M for FPPS(F96W-N127W) | FPPS-AGGGGTGGA, FPPS(F96W-N127W)-AGGGGTGGA | FPPS-(AGGGGTGGA)-NES, FPPS(F96W-N127W)-AGGGGTGGA-LS |
| AcNES1_AG4TGGA_fw | NES forward for FPPS-NES fusion protein; appends linker peptide (AGGGGTGGA) | AGGAGGTACTGGTGGTGCTGAGCTCATGGCTACCGCAGCAGGT | 43 | pJT9R | AGGGGTGGA-NES | FPPS-(AGGGGTGGA)-NES |
| ERG20_EAAAK_rv | FPPS reverse for FPPS-NES fusion protein; appends linker peptide $(EAAAK)_2$ | GGCTGCAGCTTCTTTTGCAGCAGCTTCAGATCTTTTGCTTCTCTTGTAAACTTTGTTC | 58 | pJT9R | FPPS-$(EAAAK)_2$ | FPPS-$(EAAAK)_2$-NES |
| AcNES1_EAAAK_fw | NES forward for FPPS-NES fusion protein; appends linker peptide $(EAAAK)_2$ | AAAAGAAGCTGCAGCCAAGGAGCTCATGGCTACCGCAGCAGGT | 43 | pJT9R | $(EAAAK)_2$-NES | FPPS-$(EAAAK)_2$-NES |
| ERG20_PT4P_rv | FPPS reverse for FPPS-NES fusion protein; appends linker peptide $(PT)_4P$ | GCGTTGGTGTGGGAGTAGGTGTTGGAGATCTTTTGCTTCTCTTGTAAACTTTGTTC | 56 | pJT9R | FPPS-$(PT)_4P$ | FPPS-$(PT)_4P$-NES |

| | | | | | | |
|---|---|---|---|---|---|---|
| AcNES1_PT4P_fw | NES forward for FPPS-NES fusion protein; appends linker peptide $(PT)_4P$ | TACTCCCACACCAACGCCTGAGCTCATGGCTACCGCAGCAGGT | 43 | pJT9R | $(PT)_4P$-NES | FPPS-$(PT)_4P$-NES |
| yEGFP_pGAL7_fw | GFP forward, with overhang for $P_{GAL7}$ | AAAACAGTTGAATATTCCCTCAAAAGAGCTCAAAATGGGATCCTCTAAAGGT | 52 | pILGFPB5A (Peng et al. 2015[5]) | GFP | FPPS + GFP-NES |
| yEGFP_AG4TGGA_rv | GFP reverse for GFP-NES fusion protein; appends linker peptide (AGGGGTGGA) | AGCACCACCAGTACCTCCTCCACCAGCAGATCTTTTGTACAATTCATCCATACCATG | 57 | pILGFPB5A | GFP-AGGGGTGGA | FPPS + GFP-NES |
| ClLIS1_AG4TGGA_fw | LS forward for FPPS-LS fusion protein; appends linker peptide (AGGGGTGGA) | GAGGTACTGGTGGTGCTGAGCTCATGAGAAGATCAGCTAACTATC | 45 | pJT11 (Peng et al. 2018[6]) | AGGGGTGGA-LS | FPPS(F96W-N127W)-LS |
| ClLIS1_HIS5t_rv | LS reverse, with overhang for $T_{HIS5}$ | AAACTGTACATATACTGTTTAAATTAATCTATCTCGAGCTATTAACCCTTTGTACCTGG | 59 | pJT11 | LS | FPPS(F96W-N127W) + LS, FPPS(F96W-N127W)-LS |
| ClLIS1_pGAL7_fw | LS forward (for FPPS + LS), with overhang for $P_{GAL7}$ | AAACAGTTGAATATTCCCTCAAAAGAGCTCAAAATGAGAAGATCAGCTAACTATC | 55 | pJT11 | LS | FPPS(F96W-N127W) + LS |
| PcPTS_AG4TGGA_fw | PTS forward for FPPS-PTS fusion protein; appends linker peptide (AGGGGTGGA) | GGTACTGGTGGTGCTGAGCTCATGGAATTGTATGCTCAATC | 41 | PTS gene block | AGGGGTGGA-PTS | FPPS-PTS |

| | | | | | | |
|---|---|---|---|---|---|---|
| PcPTS_HIS5t_rv | PTS reverse, with overhang for $T_{HIS5}$ | AACTGTACATATACTGTTTAAATTAATCTATCTCGAGTTAATATGGAACAGGGTGTAGG | 59 | PTS gene block | PTS | FPPS-PTS |
| Fseq_GAL10p | Forward sequencing primer; binds to $P_{GAL10}$ | GTGGTAATGCCATGTAATATGATTATTAAAC | 31 | NA | NA | All constructs |
| Rseq_KlURA3t | Reverse sequencing primer; binds to *K. lactis* URA3 terminator | GCATTGGCACGGTGCAACAC | 20 | NA | NA | All constructs |
| Rseq_AcNES1 | Reverse sequencing primer; binds close to 5’ of NES | TAAGCGTTGGAGTTTTGTGGG | 21 | NA | NA | NES fusion constructs (e.g. FPPS-NES, FPPS + GFP-NES) |
| Fseq_GAL7p | Forward sequencing primer; binds to PGAL7 | TAGTATTCGTTTGGTAAAGTAGAGG | 25 | NA | NA | Constructs with two separate proteins (e.g. FPPS + NES) |
| Rseq_GAL10t | Reverse sequencing primer; binds to TGAL10 | CTCAACAGTGCTCCGAAG | 18 | NA | NA | Constructs with two separate proteins (e.g. FPPS + NES) |
| Fscreen_127W | Colony PCR primer to screen for ERG20(F96W-N127W) variant | GGGAAATTGCCATCTGG | 17 | NA | NA | FPPS(F96W-N127W) + LS, FPPS(F96W-N127W)-LS |

**Table S2.** Synthetic gene sequences.

| **Name** | **Description** | **Sequence (5' - 3')** | **Length (bp)** |
|---|---|---|---|
| NES (or AcNES1) | *Actinidia chinensis* linalool/nerolidol synthase, codon-optimised for *S. cerevisiae*.<br><br>PCR-amplified from plasmid pJT9R (Peng et al. 2017[3]). | ATGGCTACCGCAGCAGGTCCTATCGCAACTAACAACTCCCCACAAAACTCCAACGCTTACAG<br>AACTCCAATCGCTCCTTCCGTACCAATTACTCATAAATGGTCTATAGCTGAAGATTTGACATG<br>TATTTCCAATCCTAGTAAGCACAATAACCCTCAAACTGGTTACAGATCATTTTCTGACGAATT<br>ATACGTTAAGTACGAAGAAAAGTTGGAAGATGTTAGAAAAGCATTAAGAGAAGTTGAAGAA<br>AACCCTTTGGAAGGTTTAGTTATGATAGACGCTTTGCAAAGATTGGGTATCGATTACCATTTC<br>AGAGGTGAAATTGGTGCATTCTTGCAAAAGCAACAAATCATATCTTCAACTCCAGATGGTTA<br>CCCTGAACATGGTTTGTACGAAGTTTCAACATTGTTTAGATTCTTAAGACAAGAAGGTCACA<br>ATGTTACCGCTGACGTCTTTAATAACTTCAAGGATAAGGAAGGTAGATTCAGATCAGAATTG<br>TCAACAGATATTAGAGGTTTGATGTCCTTATACGAAGCAAGTCAATTGAGAATAGAAGGTGA<br>AGACATCTTAGATCAAGCTGCTGATTTCTCCAGTCAATTGTTAGGTAGATGGACAAAAGATC<br>CTAATCATCACGAAGCCAGATTGGTTTCTAACACTTTAACACATCCATACCACAAGTCATTG<br>GCTACCTTTATGGGTCAAAAATTGTCCTACATGAACTGCAAGGGTCCAAACTGGGACGGTGT<br>CGATAATTTGCAAGAATTAGCTAAGATGGATTTGACTATCGTACAAAGTATCCATCAAAAGG<br>AAGTATTCCAAGTTTCTCAATGGTGGAAGGATACAGGTTTAGCCAATGAATTGAAATTGGCT<br>AGAAACCAACCATTGAAGTGGTATATGTGGCCTATGGCCGCTTTAACCGATCCAAGATTCAG<br>TGAAGAAAGAGTTGAATTAACTAAGCCTATTTCTTTTATATACATCATAGATGACATCTTCGA<br>CGTCTATGGTACCATTGAAGAATTGACCTTGTTTACTGATGCAGTTAATAGATGGGAATTGTC<br>TGCCGTCGAACAATTACCAGACTACATGAAAGTATGTTTCAAGGCTTTGTACGATGTTACCA<br>ACGAAATCGCATACAAAATCTATAAAAAGCATGGTCAAAACCCTATTGATTCCTTGCAAAAG<br>ACTTGGGCTAGTTTGTGCAATGCATTTTTAGTTGAAGCAAAGTGGTTCGCCTCTGGTCACTTG<br>CCAAATGCAGAAGAATACTTAAAGAACGGTATCATCTCTTCAGGTGTTCATGTTGTCTTGGC<br>CCACATGTTTTTCTTGTTAGGTGACGGTATTACACAAGAATCAGTTGATTTGGTAGATGACTA<br>TCCAGGTATTTCCACAAGTATCGCAACCATTTTGAGATTATCTGATGACTTGGGTTCAGCCAA<br>AGATGAAGACCAAGATGGTTATGATGGTTCTTACATCGAATGTTACATGAAGGAACATAAGG<br>GTTCCAGTGTCGATTCAGCCAGAGAAGAAGTAATAAGAATGATCTCCGAAGCATGGAAATG<br>TTTGAATAAGGAATGCTTATCACCAAACCCTTTTTCTGAATCATTCAGAATAGGTTCCTTGAA<br>TATGGCTAGAATGATCCCTATGATGTACTCTTACGATGACAACCATAACTTGCCAATTTTAGA<br>AGAACACATGAAGGCAATGATCTATAACACATCTTTATAA | 1722 |

| | | | |
|---|---|---|---|
| deadFPPS gene block (includes assembly overhangs for $P_{GAL7}$ and NES) | *S. cerevisiae* farnesyl diphosphate synthase (ERG20) with the K197G-K254A double point mutations. The yeast sequence also contains a G253D mutation that was inadvertently introduced during cloning.<br><br>Gene block was used directly in isothermal assembly. | GATAAAAAAAAACAGTTGAATATTCCCTCAAAAGAGCTCAAAATGGCTTCAGAAAAAGAAA<br>TTAGGAGAGAGAGATTCTTGAACGTTTTCCCTAAATTAGTAGAGGAATTGAACGCATCGCTT<br>TTGGCTTACGGTATGCCTAAGGAAGCATGTGACTGGTATGCCCACTCATTGAACTACAACAC<br>TCCAGGCGGTAAGTTAAATAGAGGTTTGTCCGTTGTGGACACGTATGCTATTCTCTCCAACA<br>AGACCGTTGAACAATTGGGGCAAGAAGAATACGAAAAGGTTGCTATTCTAGGTTGGTGCATT<br>GAGTTGTTGCAGGCTTACTTCTTGGTCGCCGATGATATGATGGACAAGTCCATTACCAGAAG<br>AGGCCAACCATGTTGGTACAAGGTTCCTGAAGTTGGGGAAATTGCCATCAATGACGCATTCA<br>TGTTAGAGGCTGCTATCTACAAGCTTTTGAAATCTCACTTCAGAAACGAAAAATACTACATA<br>GATATCACCGAATTGTTCCATGAAGTCACCTTCCAAACCGAATTGGGCCAATTGATGGACTT<br>AATCACTGCACCTGAAGACAAAGTCGACTTGAGTAAGTTCTCCCTAAAGAAGCACTCCTTCA<br>TAGTTACTTTCGGTACTGCTTACTATTCTTTCTACTTGCCTGTCGCATTGGCTATGTACGTTGC<br>CGGTATCACAGATGAAAAGGATTTGAAACAAGCCAGAGATGTCTTGATTCCATTGGGTGAAT<br>ATTTCCAAATTCAAGATGACTACTTAGACTGCTTCGGTACCCCAGAACAGATCGATGCTATC<br>GGTACAGATATCCAAGATAACAAATGTTCTTGGGTAATCAACAAGGCATTAGAACTTGCTTC<br>CGCAGAACAAAGAAAGACTTTAGACGAAAATTACGGTAAGAAGGACTCAGTCGCAGAAGCC<br>AAATGCAAAAAGATTTTCAATGACTTGAAAATCGACCAGTTATACCACGAATATGAAGAGTC<br>TGTTGCCAAGGATTTGAAGGCCAAGATCTCCCAAGTCGACGAGTCTCGTGGCTTCAAAGCCG<br>ACGTCTTAACTGCGTTTTTGAACAAAGTTTACAAGAGAAGCAAAAGATCTGCTGGTGGAGGA<br>GGTACTGGTGGTGCTGAGCTCATGGCTACCGCAGCAGGT | 1155 |
| GFP | Yeast-enhanced GFP (yeGFP3 from Cormack et al. 1997[7]).<br><br>PCR-amplified from pILGFPB5A (Peng et al. 2015[5]). | ATGGGATCCTCTAAAGGTGAAGAATTATTCACTGGTGTTGTCCCAATTTTGGTTGAATTAGAT<br>GGTGATGTTAATGGTCACAAATTTTCTGTCTCCGGTGAAGGTGAAGGTGATGCTACTTACGG<br>TAAATTGACCTTAAAATTTATTTGTACTACTGGTAAATTGCCAGTTCCATGGCCAACCTTAGT<br>CACTACTTTCGGTTATGGTGTTCAATGTTTTGCGAGATACCCAGATCATATGAAACAACATGA<br>CTTTTTCAAGTCTGCCATGCCAGAAGGTTATGTTCAAGAAAGAACTATTTTTTTCAAAGATGA<br>CGGTAACTACAAGACCAGAGCTGAAGTCAAGTTTGAAGGTGATACCTTAGTTAATAGAATCG<br>AATTAAAAGGTATTGATTTTAAAGAAGATGGTAACATTTTAGGTCACAAATTGGAATACAAC<br>TATAACTCTCACAATGTTTACATCATGGCTGACAAACAAAAGAATGGTATCAAAGTTAACTT<br>CAAAATTAGACACAACATTGAAGATGGTTCTGTTCAATTAGCTGACCATTATCAACAAAATA<br>CTCCAATTGGTGATGGTCCAGTCTTGTTACCAGACAACCATTACTTATCCACTCAATCTGCCT<br>TATCCAAAGATCCAAACGAAAAGAGAGACCACATGGTCTTGTTAGAATTTGTTACTGCTGCT<br>GGTATTACCCATGGTATGGATGAATTGTACAAAAGA | 723 |

| | | | |
|---|---|---|---|
| FPPS (F96W-N127W) | *S. cerevisiae* farnesyl diphosphate synthase (ERG20) with the F96W-N127W double point mutations.<br><br>PCR-amplified from plasmid pIT6EG7M (Peng et al. 2022[4]) for FPPS(F96W-N127W). | ATGGCTTCAGAAAAAGAAATTAGGAGAGAGAGATTCTTGAACGTTTTCCCTAAATTAGTAGA<br>GGAATTGAACGCATCGCTTTTGGCTTACGGTATGCCTAAGGAAGCATGTGACTGGTATGCCC<br>ACTCATTGAACTACAACACTCCAGGCGGTAAGCTAAATAGAGGTTTGTCCGTTGTGGACACG<br>TATGCTATTCTCTCCAACAAGACCGTTGAACAATTGGGGCAAGAAGAATACGAAAAGGTTGC<br>CATTCTAGGTTGGTGCATTGAGTTGTTGCAGGCTTACTGGTTGGTCGCCGATGATATGATGGA<br>CAAGTCCATTACCAGAAGAGGCCAACCATGTTGGTACAAGGTTCCTGAAGTTGGGGAAATTG<br>CCATCTGGGACGCATTCATGTTAGAGGCTGCTATCTACAAGCTTTTGAAATCTCACTTCAGAA<br>ACGAAAAATACTACATAGATATCACCGAATTGTTCCATGAGGTCACCTTCCAAACCGAATTG<br>GGCCAATTGATGGACTTAATCACTGCACCTGAAGACAAAGTCGACTTGAGTAAGTTCTCCCT<br>AAAGAAGCACTCCTTCATAGTTACTTTCAAGACTGCTTACTATTCTTTCTACTTGCCTGTCGC<br>ATTGGCCATGTACGTTGCCGGTATCACGGATGAAAAGGATTTGAAACAAGCCAGAGATGTCT<br>TGATTCCATTGGGTGAATACTTCCAAATTCAAGATGACTACTTAGACTGCTTCGGTACCCCAG<br>AACAGATCGGTAAGATCGGTACAGATATCCAAGATAACAAATGTTCTTGGGTAATCAACAA<br>GGCATTAGAACTTGCTTCCGCAGAACAAAGAAAGACTTTAGACGAAAATTACGGTAAGAAG<br>GACTCAGTCGCAGAAGCCAAATGCAAAAAGATTTTCAATGACTTGAAAATTGAACAGCTATA<br>CCACGAATATGAAGAGTCTATTGCCAAGGATTTGAAGGCCAAAATTTCTCAGGTCGATGAGT<br>CTCGTGGCTTCAAAGCTGATGTCTTAACTGCGTTCTTGAACAAAGTTTACAAGAGAAGCAAA | 1056 |
| LS (or ClLIS1) | *Citrus limon* limonene synthase, codon-optimised for *S. cerevisiae.*<br><br>PCR-amplified from plasmid pJT11 (Peng et al. 2018[6]). | ATGAGAAGATCAGCTAACTATCAACCATCCATTTGGGACCACGACTTTTTACAATCCTTGAA<br>CTCTAACTACACCGACGAAGCATACAAGAGAAGAGCAGAAGAATTACGTGGTAAAGTAAAG<br>ATAGCCATCAAAGATGTCATCGAACCTTTGGACCAATTGGAATTGATTGATAACTTGCAAAG<br>ATTGGGTTTAGCCCATAGATTTGAAACCGAAATCAGAAACATCTTGAACAACATCTATAACA<br>ACAATAAGGATTACAACTGGAGAAAGGAAAATTTGTACGCTACTTCCTTGGAATTCAGATTG<br>TTAAGACAACACGGTTACCCAGTCAGTCAAGAAGTTTTTAACGGTTTCAAGGATGACCAAGG<br>TGGTTTTATTTGTGATGACTTCAAGGGTATATTGTCCTTGCATGAAGCTTCTTACTACTCATTG<br>GAAGGTGAAAGTATAATGGAAGAAGCATGGCAATTCACTTCCAAACACTTGAAGGAAGTTA<br>TGATAAGTAAAAATATGGAAGAAGATGTTTTCGTAGCTGAACAAGCAAAGAGAGCCTTGGA<br>ATTACCATTGCATTGGAAAGTTCCTATGTTGGAAGCAAGATGGTTCATCCATATATACGAAA<br>GAAGAGAAGATAAGAACCACTTGTTGTTGGAATTGGCAAAGATGGAATTCAATACATTACA<br>AGCCATCTATCAAGAAGAATTGAAGGAAATCTCTGGTTGGTGGAAGGATACCGGTTTGGGTG<br>AAAAGTTGTCATTCGCTAGAAATAGATTGGTCGCATCTTTCTTATGGTCAATGGGTATTGCCT<br>TTGAACCTCAATTCGCTTACTGTAGAAGAGTTTTGACAATCTCTATTGCATTGATCACCGTTA<br>TAGATGACATCTATGACGTATACGGTACTTTGGATGAATTGGAAATCTTTACAGACGCTGTT | 1671 |

| | | | |
|---|---|---|---|
| | | GAAAGATGGGATATCAACTATGCATTAAAGCATTTGCCAGGTTACATGAAGATGTGCTTTTT<br>AGCCTTGTACAACTTCGTCAACGAATTCGCTTACTACGTTTTGAAGCAACAAGATTTCGACTT<br>ATTGTTATCTATTAAAAACGCTTGGTTGGGTTTGATACAAGCCTATTTGGTAGAGGCTAAGTG<br>GTACCATTCTAAGTACACACCTAAGTTGGAAGAATACTTGGAAAACGGTTTAGTCTCAATCA<br>CTGGTCCATTGATCATCACAATCTCCTATTTGAGTGGTACTAACCCTATCATTAAAAAGGAAT<br>TGGAATTCTTGGAATCAAACCCAGATATCGTTCACTGGTCTTCAAAAATTTTCAGATTGCAAG<br>ATGACTTAGGTACTTCCAGTGACGAAATTCAAAGAGGTGACGTACCTAAATCTATACAATGC<br>TACATGCATGAAACAGGTGCATCAGAAGAAGTTGCCAGACAACACATTAAGGATATGATGA<br>GACAAATGTGGAAAAAGGTTAATGCTTACACCGCAGATAAAGACTCCCCATTGACTGGTACT<br>ACAACCGAATTCTTGTTGAACTTAGTTAGAATGTCTCATTTCATGTATTTGCATGGTGACGGT<br>CACGGTGTACAAAATCAAGAAACCATTGATGTCGGTTTTACTTTGTTATTCCAACCTATTCCA<br>TTAGAAGATAAACACATGGCATTCACCGCATCACCAGGTACAAAGGGTTAATAG | |
| PTS (or PcPTS) gene block | *Pogostemon cablin* patchoulol synthase, codon-optimised for *S. cerevisiae.* PTS gene is flanked by SacI and XhoI restriction enzyme sites.<br><br>Gene block was used directly in isothermal assembly for cloning FPPS + PTS, and includes assembly overhangs for $P_{GAL7}$ and $T_{HIS5}$. | GATAAAAAAAAACAGTTGAATATTCCCTCAAAAGAGCTCAAAATGGAATTGTATGCTCAATC<br>TGTTGGTGTTGGTGCTGCATCAAGACCTCTAGCCAATTTTCATCAATGTGTATGGGGAGACA<br>AATTCATTGTTTACAACCCACAATCATCCCAGGCTGGAGAGAGAGAACAGGCTGAGGAGCTT<br>AAAGTAGAGCTAAAAAGAGAGCTTAAGGAAGCATCAGATAATTATATGAGGCAACTGAAAA<br>TGGTAGATGCAATACAACGTTTAGGCATTGACTATTTATTTGTCGAAGATGTTGATGAAGCTT<br>TGAAGAATTTGTTTGAAATGTTTGATGCTTTCTGCAAGAATAATCATGACATGCACGCCACTG<br>CTCTATCCTTTCGTCTTCTAAGACAACATGGCTATAGAGTTTCATGTGAAGTTTTTGAAAAGT<br>TTAAGGATGGCAAAGATGGATTTAAGGTTCCAAATGAGGATGGAGCCGTTGCAGTCCTTGAA<br>TTCTTCGAAGCCACTCATTTGAGAGTCCATGGAGAAGACGTCCTTGATAATGCTTTTGTCTTC<br>ACTAGGAACTACTTGGAATCAGTCTATGCAACTTTGAACGATCCAACCGCGAATCAAGTCCA<br>CAACGCATTGAATGAGTTCTCTTTTAGGAGAGGATTGCCACGTGTGGAAGCAAGGAAGTACA<br>TATCAATCTACGAGCAATACGCATCTCATCACAAAGGCTTGTTAAAACTTGCTAAGCTGGAT<br>TTCAACTTGGTACAAGCTTTGCACAGAAGGGAGCTGAGTGAAGATTCTAGGTGGTGGAAGAC<br>TTTACAAGTGCCCACAGAGCTATCATTCGTTAGAGATAGGTTGGTGGAGTCCTACTTTTGGGC<br>TTCGGGATCTTATTTCGAACCGAATTATTCGGTAGCTAGGATGATTTTAGCAAAAGGATTAG<br>CTGTATTATCTCTTATGGACGATGTGTATGATGCATATGGTACTTTTGAGGAATTACAAGTGT<br>TCACAGATGCAATCGAAAGGTGGGATGCTTCATGTTTAGATAAACTTCCAGAGTACATGAAA<br>ATAGTATACAAGGCCCTTTTGGATGTGTTTGAGGAAGTTGACGAGGAGGTGATCAAGCTAGG<br>TGCACCATATAGGGTCTACTATGGAAAAGAAGCCATGAAATATGCCGCTAGAGCTTACATGG<br>AAGAGGCCCAATGGAGGGAGCAAAAGCACAAACCCACAACTAAGGAGTATATGAAGCTGG | 1738 |

| | | |
|---|---|---|
| | | CAACAAAGACCTGTGGCTATATAACTCTAATAATATTATCATTTCTTGGAGTGGAAGAGGGC<br>ATTGTGACTAAAGAAGCCTTCGATTGGGTTTTCTCCAGACCTCCTTTCGTCGAGGCTACTTTA<br>ATCATTGCTAGGTTGATCAATGATATTACAGGATGTGAATTTGAGAATAAAAGGGAGCACGT<br>TAGAACTGCAGTCGAATGCTACATGGAAGAGCACAAAGTGGGAAAGCAAGAGGTGGTCTCT<br>GAATTTTACAACCAAATGGAGTCAGCATGGAAGGACATAAATGAGTGCTTGCTAAGACCAG<br>CTGAATTTCCAATCCCTCTACTTAATCTTATTTTAAATTCTGTCAGAACACTTGAGGTTATTTA<br>CAAAGAGGGCGATTCCTATACACACGTGGGTCCTGCAATGCAAAACATCATCAAGCAGTTGT<br>ACCTACACCCTGTTCCATATTAACTCGAGATAGATTAATTTAAACAGTATATGTACAGTT |

**Table S3.** Comparing the relative levels of terpene synthase and the corresponding isoprenoid product.

Values shown are the mean of 3 biological replicates. 'N/A' indicates strains that were not characterised by proteomics. Dry cell mass values were estimated by multiplying the $OD_{600}$ reading at 72 h by a published conversion factor (0.644 g DCM/L per unit $OD_{600}$; ref. Myers et al. 2013[1]). The fold changes presented here are calculated based on DCM-normalised titres.

| | **FPPS fold change** | **NES fold change** | **Nerolidol (mg/L)** | **Nerolidol (mg/g DCM)** | **Nerolidol fold change** |
|---|---|---|---|---|---|
| FPPS + NES | 1.00 | 1.00 | 29.65 | 2.26 | 1.00 |
| FPPS-NES (AGGGGTGGA) | 1.40 | 28.47 | 3509.76 | 197.05 | 87.06 |
| FPPS-NES $(AGGGGTGGA)_2$ | N/A | N/A | 4204.13 | 230.78 | 101.96 |
| FPPS-NES $(EAAAK)_2$ | N/A | N/A | 4071.04 | 223.11 | 98.57 |
| FPPS-NES $(PT)_4P$ | N/A | N/A | 3561.54 | 193.18 | 85.35 |
| FPPS + deadFPPS-NES | 3.81 | 23.40 | 3029.67 | 176.42 | 77.94 |
| FPPS + GFP-NES | 3.21 | 16.53 | 2985.83 | 193.15 | 85.33 |
| | **FPPS fold change** | **LS fold change** | **Limonene (mg/L)** | **Limonene (mg/g DCM)** | **Limonene fold change** |
| FPPS(F96W-N127W) + LS | 1.00 | 1.00 | 72.28 | 5.69 | 1.00 |
| FPPS(F96W-N127W)-LS | 0.53 | 1.48 | 134.80 | 6.61 | 1.16 |
| | **FPPS fold change** | **PTS fold change** | **Patchoulol (mg/L)** | **Patchoulol (mg/g DCM)** | **Patchoulol fold change** |
| FPPS + PTS | 1.00 | 1.00 | 0.51 | 0.049 | 1.00 |
| FPPS-PTS | 0.41 | 4.03 | 1.97 | 0.192 | 3.95 |
